# Structured dynamics of neural activity across developing neocortex

**DOI:** 10.1101/012237

**Authors:** James B. Ackman, Hongkui Zeng, Michael C. Crair

## Abstract

The cerebral cortex exhibits spontaneous and sensory evoked patterns of activity during early development that is vital for the formation and refinement of neural circuits. Identifying the source and flow of this activity locally and globally is critical for understanding principles guiding self-organization in the developing brain. Here we use whole brain transcranial optical imaging at high spatial and temporal resolution to demonstrate that dynamical patterns of neuronal activity in developing mouse neocortex consist of spatially discrete domains that are coordinated in an age, areal, and behavior-dependent fashion. Ongoing cortical activity displays mirror-symmetric activation patterns across the cerebral hemispheres and stereotyped network architectures that are shaped during development, with parietal-sensorimotor subnetworks functionally connected to occipital regions through frontal-medial cortical areas. This study provides the first broad description of population activity in the developing neocortex at a scope and scale that bridges the microscopic and macroscopic spatiotemporal resolutions provided by traditional neurophysiological and functional neuroimaging techniques. Mesoscale maps of cortical population dynamics within animal models will be crucial for future efforts to understand and treat neurodevelopmental disorders.

## Introduction

Neuronal activity is required for the proper morphologic and functional development of the vertebrate brain^1,2^. Early in gestation, calcium transients associated with activity modulate neuronal proliferation, migration, differentiation, and neurotransmitter specification^1,3^. In mid-gestation, spontaneous motor neuron activity and associated embryonic limb movements shape neuromuscular junction formation and the self-organization of spinal cord circuitry in a wide variety of species^4,5,6^. Still later in development, spontaneous and sensory driven activity are vital for the formation and refinement of neural circuits in the spinal cord, brainstem and cortex^2,7,8^. In humans, disrupting neurotransmission during fetal development results in severe brain malformations, epilepsy and cognitive disorders, underscoring the importance of activity for normal brain development^9,10^.

It is only recently, however, that a better appreciation is emerging of the complex spatial and temporal patterns of persistent neural activity that exist in the perinatal brain *in vivo.* For example, sensory-motor feedback associated with spontaneous movement generated by spinal motor neurons triggers synchronized ‘spindle-burst’ potentials among cells in somatosensory cortex before the start of locomotion and tactile behavior^11,12^. EEG recordings in humans also demonstrate spindle-burst oscillations and slow activity transients in the somatosensory and occipital cortices before birth^13,14^. Correlated bursts of activity in the developing rat hippocampus *in vivo*^15,16^, and spontaneous retinal waves that drive patterned activation of circuits throughout the visual system before the onset of vision^17,18^ are further examples of complex spatiotemporal patterned activity in the neonatal CNS *in vivo.* However, because we lack methods to assess neuronal activity at adequate temporal and spatial resolution, a holistic account of the dynamical patterns of persistent activity across the neocortex throughout neonatal development *in vivo* has never been undertaken. Here, we use genetically encoded calcium indicators and transcranial imaging to examine the emergence and pattern of spontaneous activity throughout the neonatal mouse neocortex at unprecedented spatial and temporal resolution *in vivo*. This work provides the first report on the dynamical nature of cortical activity across the developing cerebral hemispheres and its distinct and integrative features among brain regions during formation of cortical network architecture.

## Results

### Ongoing neocortical activity characterized by discrete domains

We performed transcranial optical recordings from mice expressing the genetically encoded calcium reporter GCaMP (GCaMP3 or GCaMP6) in cortical neurons to assess neuronal population activity patterns at macroscopic scale (across the entire neocortex) with mesoscopic spatial (10s of microns) and temporal resolution (100s of milliseconds). We performed this functional mesoscopic optical imaging (fMOI) in three age groups through the first two postnatal weeks, during which the mouse brain attains >90% of its adult weight^19^: P2–P5 (N = 6 mice), P8–P9 (N = 4), and P12–13 (N = 5).

The neocortex exhibits a characteristic modular organization across the cortical surface in which vertical arrays of cells, derived from the same or nearby embryonic precursors, are grouped together as functional columns^20^. Cortical columns range from 300–600 μm in diameter, even across species whose brain volume differs by greater than a factor of one-thousand^20^. fMOI revealed that supracellular cortical activity patterns were characterized by discontinuous, discrete domains (**Fig. 1a-c**) (**Supplementary Movie 1**). These domains were active from 0.4 – 2.6 s at a time (10-90th percentiles) and ranged from 250 – 976 μm in diameter (**Fig. 1e-h**) (**Table 1**). The duration of cortical domain activity was not significantly affected by age (F = 0.933, p = 0.428) (P2–5, N = 15653; P8–9, N = 70189; P12–13, N = 120214 domains) (**Fig. 1e,f**), but the diameter (F = 25.788, p = 0.000188), (**Fig. 1g,h**), and the frequency of cortical domain activations increased significantly with age (F = 29.562, p = 8.86e–12) (P2–5, N = 22; P8–9, N = 30; P12–13, N = 38 movies/hemi) (**Fig. i,j**) (**Table 1**) (**Supplementary Movie 2, 3**). None of these measures differed significantly between the two hemispheres (duration: F = 0.017, p = 0.900; diameter: F = 0.192, p = 0.671808; frequency: F = 0.012, p = 0.911).

**Figure 1.**
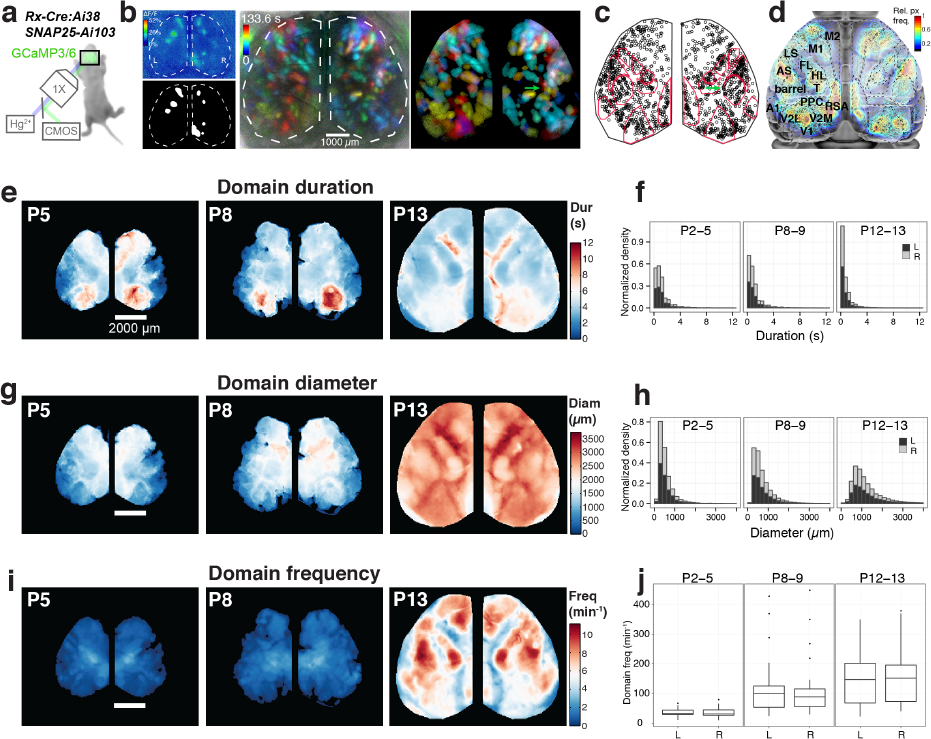
Calcium domains throughout neonatal mouse neocortex. **a**, Experimental schematic. **b**, Left panel: Single image frame showing calcium domains in both hemispheres at postnatal day 3 (P3) and the mask of detected domain signals. Middle and right panels: Time projection map from a raw dF/F movie segment and the corresponding map from automatically detected domain masks. Notice the individual domains of activity in the area of barrel cortex (arrow) **c**, Centroid positions for segmented domain masks from a 10 min recording. Points are overlaid on a reference map of primary sensory areas determined by thalamocortical inputs (red outlines). Notice rows of whisker barrels are evident in the structure of domain centroid positions (arrow). **d**, Functional activity map at P3. Based on pixel activation frequency from all detected domains in a single 10 min recording. Map is overlaid on cortical areal parcellations. Notice localized maxima and minima of functional activity between areas that approximate known anatomical cortical area boundaries and the mirroring of map structure bilaterally. **e**, Mean domain duration maps from 3 SNAP25-Ai103 mice. **f**, Histograms showing domain durations distributions in the P2–5, P8–9, and P12–13 age groups and by cortical hemisphere (L, R). **g**, Mean domain diameter maps from same 3 mice in e. **h**, Histograms showing the distributions of domain diameters. **i**, Mean domain frequency maps from same 3 mice in e. **j**, Boxplot distributions of hemispheric domain frequencies.

**Table 1.** Domain statistics.

|  | **duration (s)** | **diameter ($\mu\text{m}$)** | **frequency**<br>**(hemisphere-min<sup>-1</sup>)** |
| --- | --- | --- | --- |
| P2–5 | 0.8 (0.4) | 441.12 (147.04) | 30.90 (9.55) |
| P8–9 | 0.6 (0.2) | 569.78 (220.56) | 98.35 (27.60) |
| P12–13 | 0.4 (0.2) | 1047.66 (367.60) | 147.80 (78.65) |

Notes: Values are reported as medians (median absolute deviation)

It’s notable that the size of the active cortical domains is in close agreement with previous work showing that population activity in neonatal rat barrel cortex is synchronized among local groups of neurons and maps onto ontogenetic modules centered on each barrel column^21,22,23^. Indeed, we found a cortical area in primary somatosensory cortex at P2–5 where cortical domain activations grouped into rows and individual modules that match primary barrel cortex structure (Fig. 1c) (**Supplementary Fig. 1**). Since barrels are an archetypal model for cortical columns in the rodent, these results indicate that early domain activity across the developing cortex is related to the functional columns thought to be a fundamental processing unit of the cerebral cortex.

### Cortical dynamics differs with area and age

We examined how the spatiotemporal properties of cortical domains vary among different cortical areas by parcellating the brain into distinct anatomical regions using reference coordinates from a mouse line that expressed a tdtomato reporter in thalamocortical afferents at P7 (Fig. 1c,d, **Supplementary Fig. 1**). Patterns of thalamocortical axon terminals outline primary sensory cortical areas^24^ during mouse postnatal development. We aligned these parcellations to the Allen brain mouse atlas and then scaled the resulting cortical area reference coordinates to match activity maps from each animal containing functional boundaries for barrel cortex and visual cortex where spontaneous retinal waves functionally map out developing visual areas^17^ (Fig. 1c-e,g,i).

Cortical domain frequency among different regions scaled as a function of net cortical area, and this association became stronger during the course of development (Fig. 2a). The most frequently active cortical regions at each age group when normalized to the total amount of cortical space were the limb/trunk representations in somatosensory cortex (Fig. 1i, **Supplementary Fig. 2**). In contrast to diameter and duration, the normalized frequency and amplitude of cortical domain activity was remarkably uniform across areas at each age of development (**Supplementary Fig. 2**), suggesting a homeostatic regulation of global activity levels. The long tails in the domain duration and diameter distributions at P2–5 and P8–9 (Fig. 1f,h) were largely due to retinal wave driven cortical activity in V1 (P2–5, 37.3%; P8–9, 17.7% of all cortical domains > 5s) and wave-like activations in motor cortex (Fig. 1e, Fig. 2b,c) (P2–5, 23.7%; P8–9, 21.8% of all domains > 5s). In contrast, the next largest areal contribution to long-lasting cortical domains (durations > 5s), from V1L and barrel cortex, was considerably smaller (P2–5: V1L, 5.6%, barrel 1.2%; P8–9: V1L, 6.7%, barrel, 7.2%).

**Figure 2.**
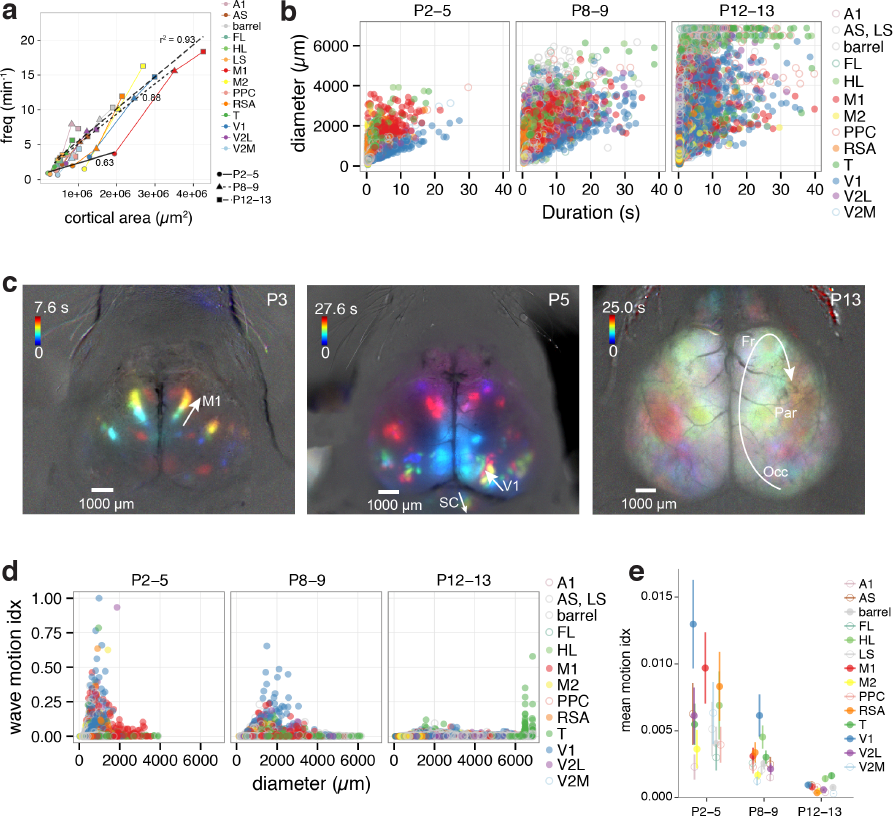
Spatiotemporal properties of cortical domains. **a**, Domain frequency as function of cortical area size. **b**, Scatterplots of domain diameter and duration. **c**, Time projection maps of waves in motor cortex at P3, visual cortex at P5, and occipital-parietal-frontal cortex at P13. **d**, Scatterplots of wave motion index as function of domain diameter. **e**, Mean wave motion index over development.

To further characterize domain activations across development and cortical regions, we computed a wave motion index, which varied significantly with age and brain area (age: F = 104.068, p = < 2e–16; area: F = 3.848, p = 1.88e–06, two-way ANOVA). Indeed, the cortical regions with the highest wave motion indices were V1 and M1 at P2–5, with V1 continuing to have the highest index at P8–9 and then dropping to a mean motion index level similar to other cortical regions at P12–13 (Fig. 2d,e). At P12–13, wave motion was much less common in all areas, but interestingly a small subpopulation of long-lasting activations (57.5 ± 11.4 s vs 0.4 ± 0.2 for all other domains) had a high wave motion index (0.055 ± 0.038 vs 1.006e–04 ± 8.606e–05), with diameters approaching that of the entire hemisphere (6488.14 ± 18.38 μm vs 1029.28 ± 367.6 μm), but occurring very infrequently (0.033% of all domains, 13.2% of all active pixels at P12–13, 0.1 events per hemisphere-min) (Fig. 2b-e) (**Supplementary Movie 4**). These global population events at P12–13 synchronized activity across cortical areas and had centers of mass that were concentrated near the middle of each hemisphere in the S1-limb/body area (Fig. 2d). These results show that the spatiotemporal properties of ongoing cortical activity is regulated in an age and areal dependent fashion and that differences in dynamical activity patterns may reflect unique developmental requirements for functional circuits locally and globally among brain regions.

### Cortical activity is coordinated with motor behavior

We monitored animal movements simultaneously with cortical activity during our fMOI recordings to gain insight into the relationship between motor activity, behavioral state and cerebral cortical dynamics during development. General anesthesia is known to abolish spontaneous retinal wave activity in the visual system^17,18^ as well as spontaneous activity in entorhinal cortex^25^. We found that activity across the entire cortex rapidly decreased upon the administration of isoflurane anesthesia (Fig. 3b,c) (**Supplementary Movie 5**) at all ages tested (P2–5, N = 2; P8–9, N = 4; P12–13, N = 3 animals). In young neonates (P2–P9), little cortical activity was observed within 20 min of deep anesthesia onset (0.37 ± 0.22 domains/hemisphere-min), while at P12–13 there was altered spontaneous activity patterns, with short duration, large diameter activations synchronizing multiple cortical regions (1.43 ± 0.41 domains/hemisphere-min; p = 0.00821 Wilcoxon Rank Sum test vs P2–9) (**Supplementary Movie 6**). Despite the rapid cessation of all cortical activity under anesthesia in young neonates, motor twitches continued (Fig. 3b), presumably through the autonomous activity of spinal motor neurons^26^. The continued spinal motor activity during early anesthesia and the altered cortical activity patterns that ensue under anesthesia at P12–13 suggests a maturational dependence of general anesthesia on network activity that affects brain regions differentially during development. Furthermore, spontaneous slow oscillations in corticothalamic networks begin during sleep and general anesthesia in rodent and cat near the end of the second postnatal week^27,28,29^—therefore it is likely that persistent cortical activity under deep anesthesia at P12–13 reflects lasting recurrent excitatory connectivity characteristic of more mature corticothalamic networks.

**Figure 3.**
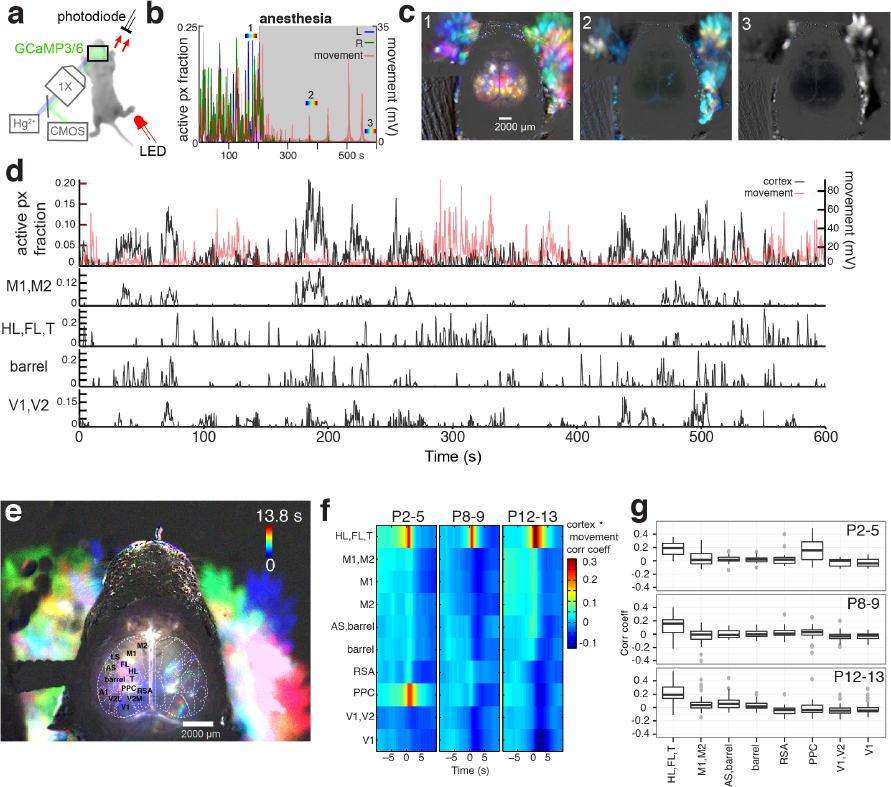
Cortical domain activity is state dependent. **a**, Experimental schematic. Red light illumination measured with a photodiode was used to monitor motor movements. **b**, Cortical activity (active pixel fraction by hemisphere) and motor movement signal after onset of isoflurane anesthesia at 205 s. **c**, Time projection maps (40 s segments) at times indicated in recording from b. **d**, Cortical activity and coincident motor movement activity signals. Active pixel fraction traces for motor (M1, M2), somatosensory (HL, FL, T; barrel), and visual (V1, V2) cortex shown at bottom of panel. **e**, Time projection map at P5 showing HL, FL, T and PPC activity coincident with motor movements. **f**, Mean cross-correlation functions between cortical regions and motor movement signals across all movies. Notice the high positive correlation between motor movement and S1-limb/body (HL, FL, T) signals at all ages. **g**, Boxplots of cortical activity and motor movement correlation at lag zero.

In the unanaesthetized animal, the relationship between motor movements and cortical activity varied depending on brain area (F = 33.7975, p < 2.2e–16) and age (F = 11.1671, p = 1.627e–05, two-way ANOVA) (P2–5, N = 22; P8–9, N = 30; P12–13, N = 38 movies) (Fig 3d-f). At all ages examined (P2–5, P8–9 and P12–13), activity in sensory cortex (S1-limb/body) was strongly correlated with movement, consistent with movement driven somatosensory self-stimulation^30^. At P2–5, activity in PPC was also strongly correlated with animal movement (Fig. 3e,f) but not at P12–13, suggesting the presence of transient input to PPC from the somatosensory periphery. Interestingly, the first age group in which activity in motor cortex exhibited significant positive correlation with motor movements was at P12–13 (r = 0.062 ± 0.019, p = 0.001449, t-test) (Fig. 3e,f). Before this (P2–5 and P8–9), motor cortex activity was not correlated with animal movements. These results indicate that spontaneous motor twitches, which are typical during perinatal development^5,31,32,33,34^, occur independent of motor cortex activity and anesthesia, but are powerful drivers of activity in somatosensory cortex in the unanaesthetized animal. Moreover, at around P12–13—a period of time when patterned vision, hearing, and locomotor exploration is beginning in mice— there is a shift towards positive preceding or zero lag correlation between frontal cortex activity and motor movements, perhaps coinciding with the beginning of goal directed behavior.

### Cortical activity comprises distinct subnetworks

The complex dynamical activity apparent in our fMOI recordings of nascent neocortical networks may reflect the emergence of functional circuitry, and could play a fundamental role in the development of intra- and inter-hemispheric connectivity^35^. To better understand the early activity patterns that may regulate cortical development and function, we examined activity correlations within and between the two hemispheres. Notably, mean activity in each hemisphere exhibited high temporal and spatial correlations across the midline (temporal, r = 0.59 ± 0.02; anterior-posterior, r = 0.43 ± 0.02; medial-lateral, r = 0.43 ± 0.02; N = 90 movies), with the strength of these inter-hemispheric correlations changing with age (temporal: F = 7.801, p = 0.000765; anterior-posterior: F = 14.63, p = 3.34e–06; medial-lateral: F = 3.663, p = 0.0297; one-way ANOVA) (**Supplementary Fig. 3**). These correlations are readily apparent in still frames (**Supplementary Fig. 3**) and movies (**Supplementary Movies 1–3**), which show frequent episodes of mirror symmetric activity between the two hemispheres across the midline.

For a more detailed analysis of the activity patterns, we computed a pairwise correlation coefficient of activity between all cortical areal parcellations in the two hemispheres (Fig. 4a). The matrix of activity correlations amongst all areas within and between the two hemispheres reveals strong, modest, and even negative correlations between areas. To examine the community structure among the cortical areas or ‘nodes’ in the pairwise association matrix, we used a hierarchical clustering algorithm based on the optimization of a graph modularity score, which measures the number of edges that fall within groups minus the number expected by chance^36^. The resulting community dendrogram (at P12–13) was used to order the mean correlation matrix for all age groups (Fig. 4a) (P2–5, N = 22/6; P8–9, N = 30/4; P12–13, N = 38/5 movies/mice). This analysis revealed a network organization and a coarse correlational structure that was similar between animals and across development (Fig. 4a). There were 4 primary network modules detected at P12–13; motor-S1-face, S1-body, medial–visual, and auditory. Indeed the 4 network modules detected at P12–13 are obvious as clusters along the diagonal in the correlation matrix, which were also apparent in the earlier age groups. Remarkably, detection of community structure in each age group independently showed similar node memberships in the 3 largest modules, with developmental switches in module membership occurring for areas PPC and M2, two brain regions known to be integral for multimodal sensory processing and motor planning (Fig. 4b).

**Figure 4.**
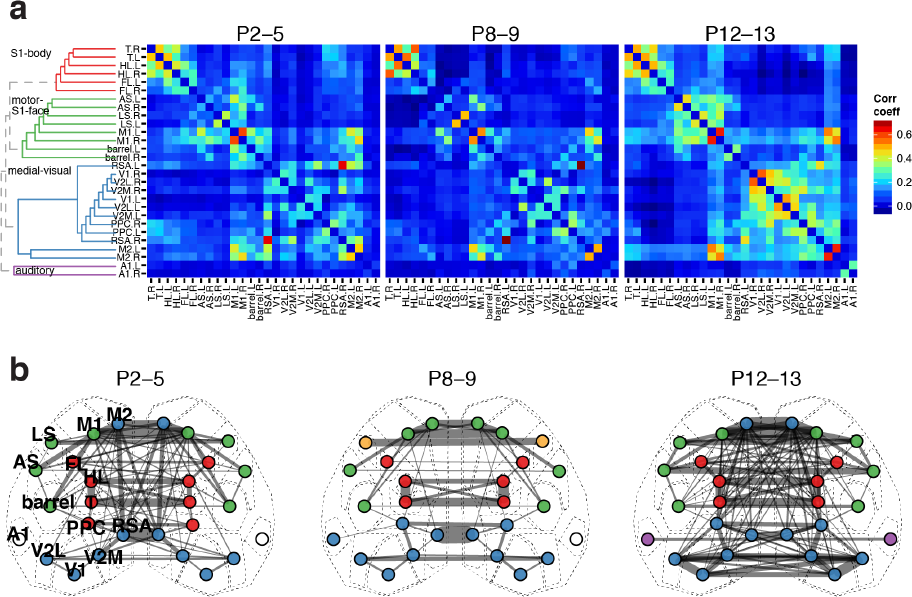
Functional architecture of developing neocortex. **a**, Group averaged correlation matrices of domain activity among cortical areas. Color map indicates Pearson’s r correlation coefficient values. Dendrogram and node order from community structure detected with hierarchical clustering in the P12–13 group. **b**, Map of cortical area associations for r > 0.15. Node colors represent cortical communities detected with hierarchical clustering within each age group. Edge width indicates the squared connection weight (r^2^). Note both similarities in module membership and increased connection strength with age.

Functional connectivity between cortical regions was highly dynamic across development, with both increases and decreases in correlation depending on brain area and age. The strongest correlations at each age typically occurred at symmetrical regions between the hemispheres in RSA, M1, M2, and T (Fig. 4a,b). In primary sensory areas such as A1, V1, and S1-barrel cortex, inter-hemispheric correlation was initially weak, but became strong at P12–13. At P8–9, correlations among cortical regions were more spread out relative to P2–5, with strong correlations becoming stronger and weak correlations becoming weaker. At P12–13, there were increased positive correlations among areas within modules and stronger negative correlations between modules. For example, visual regions exhibited increased negative correlations with the body-parietal subnetwork, A1 showed increased negative correlations with most regions, and motor-parietal areas exhibited increased intra-module correlations.

We analyzed the topological properties of developing mouse cortical networks with graph theoretic measures used in fMRI studies of large-scale functional connectivity^37^. Graphs of mean functional connectivity illustrated decreased randomness and tighter clustering among cortical modules during network development (Fig. 5a). The average path length – the mean shortest path between all node pairs – changed little over the course of development (Fig. 5c) (F = 3.153, p = 0.04765, one-way ANOVA) and did not differ significantly from equivalent random networks at any age group. In contrast, the global clustering coefficient– reflecting how highly interconnected each node’s neighbors are– significantly increased during development (Fig. 5c) (F = 18.491, p = 2.033e–07, one-way ANOVA) and was significantly greater than that of equivalent random networks at all ages. The mean degree (number of edges per node) was higher at P12–13 (two-way ANOVA: F = 33.83, p = 5.81e–08; Tukey HSD: P2–5:P8–9, p = 0.0106535; P2–5:P12–13, p = 0.0000983, P8–9:P12–13, p = 0) (Supplementary Fig. 4) and varied significantly by area (Fig. 5b) (F = 10.30, p = 3.51e–07). Mean node strength–the sum of all edge weights per node– was also significantly affected by both age and area (two-way ANOVA: age, F = 27.477, p = 3.87e–07; node, F = 8.957, p = 1.39e–06) and was highest at P12–13 (Tukey HSD: P2–5:P8–9, p = 0.0105873; P2–5:P12–13, p = 0.0007400, P8–9:P12–13, p = 0.0000002). The five cortical areas having the highest degree and node strength at all ages were M1, M2, PPC, V2M, and RSA (Fig. 5b).

**Figure 5.**
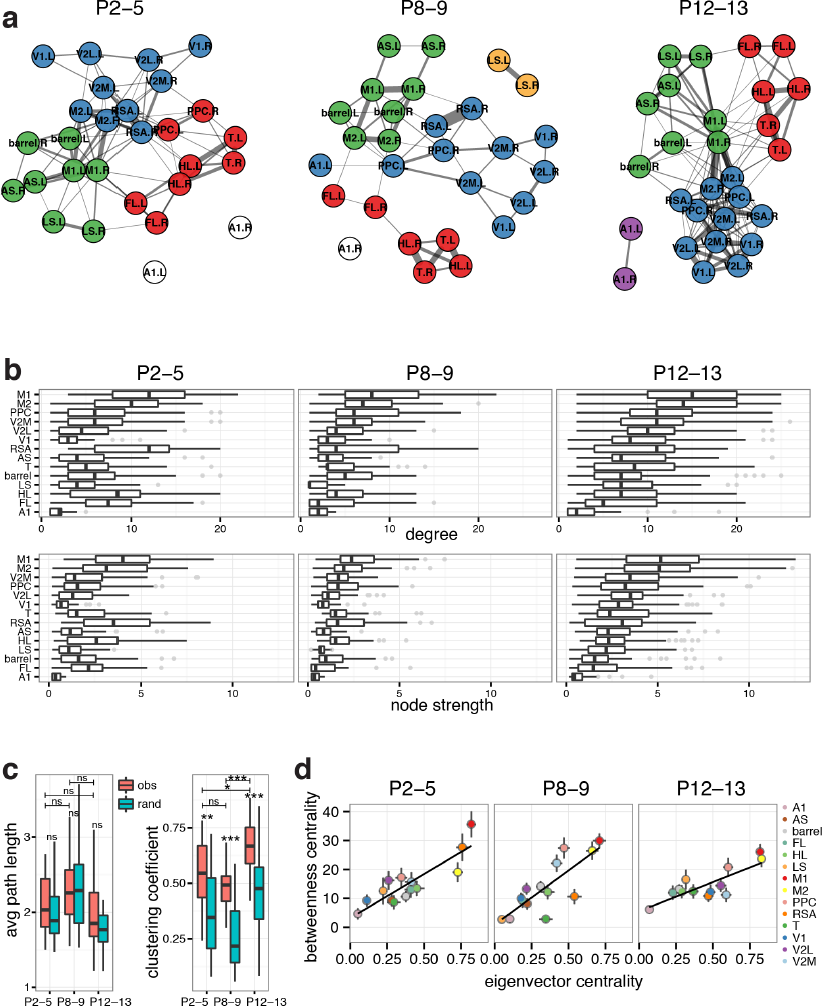
Dynamics of functional connectivity in developing neocortex. **a**, Graph of functional connections for r > 0.15. Node colors represent cortical communities detected with hierarchical clustering within each age group. **b**, Boxplots of degree (number of links) and node strength (sum of connection weights) by cortical area. The distributions become increasingly ordered like the P12–13 group with age. **c**, Boxplots of clustering coefficient and average path length by recording. **d**, Scatterplots of mean network centrality scores by cortical area. Error bars are s.e.m.

We computed measures of node centrality to identify potential hubs in the cortical network. Betweenness centrality identifies important throughputs in a network by measuring the fraction of all shortest paths that pass through a node^38^. Betweenness centrality varied significantly by node but not with age (node: F = 17.320, p < 2e–16; age: F = 0.869, p = 0.42; two-way ANOVA) (Fig. 5d). In contrast, eigenvector centrality—which is proportional to the sum of centralities for a node’s connections, giving larger scores to nodes that are linked to highly connected nodes—varied significantly by node but also increased with age (node: F = 88.29, p < 2e–16; age: F = 6.84, p < 2e–16; two-way ANOVA) (Fig. 5d). High scores in both centrality measures indicate hub nodes^38^, which included M1 and M2. Deviations from betweenness-eigenvector linearity can indicate whether a node is potentially more important as a network throughput (higher betweenness centrality) or as a driver node (higher eigenvector centrality). Interestingly, V1 and V2L shifted towards larger eigenvector centrality at P12–13, but maintained low betweenness centrality indicating that visual cortex may be positioned to have greater influence on network activity around the time of eye opening through action as a driver node. These results indicate that greater local connectivity together with higher connection strengths are key features in the development of global network architecture of the cerebral cortex. Furthermore, the connection of occipital to parietal-sensorimotor network modules through frontal-medial cortical hubs such as M2/cingulate, RSA, and posterior parietal cortex suggests that the developing mouse cerebral cortex shares key features of network topology observed in rat, primate, and human resting state brain networks.

## Discussion

We have provided the first account of neuronal population activity at high spatial and temporal resolution across the developing cortex *in vivo*. Characteristic features of brain activity varied with age and with cortical region. At early postnatal ages (P2–5), activity in some cortical areas was wave-like, particularly the primary visual (V1) and motor (M1) cortices, consistent with earlier reports of waves in V1 originating in the retina^8^. Wave-like activity persisted in V1 until P8–9, but elsewhere the activity domains were generally more static and became more frequent with age. In contrast to classical *in vitro* calcium imaging work^39^, but similar to more recent functional imaging and electrophysiology studies in rodent^21,22,23^, ongoing cortical domain activity in the neonatal barrel cortex was associated with functional columns. Interestingly, domain activity throughout the neocortex increased in diameter by a factor of 2–3x during the course of the second postnatal week– an amount greater than the 1.5x increase in linear growth along the medial-lateral or anterior-posterior surface of the mouse necortex from P3 to P13. Notably, there was spontaneous domain activity in primary auditory cortex (A1) at all ages studied, even before the end of the second postnatal week when hearing begins in mice. This indicates the presence of spontaneous activity in the auditory system, likely originating in the cochlea^40^, that may coordinate patterned activity in the developing auditory system, much as spontaneous retinal waves do in the visual system before onset of vision^17^.

Consistent with reports of twitch-activated local field potentials (‘spindle-bursts’) in S1 and M1 during development^12,30,41^, cortical activity was coordinated with motor movements depending upon area and age. The high spatial resolution of our fMOI recordings showed that transient cortical domain activity was positively associated with motor movements primarily in somatosensory cortex and posterior parietal cortex (PPC) in young neonates. Throughout somatosensory cortex, activity was closely correlated with limb, body, and facial movements that generate sensory stimulation^30^. Remarkably, activity in motor cortex was not associated with general limb and body movements until P12–13, presumably because movement in young neonates is initiated in the brainstem and spinal cord, not motor cortex^5,26,30^, suggesting that ongoing activity in frontal-motor cortex might be linked to the development of higher order associative connections within the cerebral cortex. The PPC is a multimodal sensorimotor processing region important for perceptual decision making, attention, and movement planning in primates and memory-dependent spatial navigation and decision tasks in rodents^42,43^. The PPC receives input from sensory cortical areas, association thalamic nuclei (LD, LP), and retrosplenial cortex (RSA) and may influence motor planning in frontal cortex via its major axonal output to prefrontal cortex, secondary motor cortex (M2), striatum, superior colliculus, and LD/LP^43^. Our observation of an age dependent correlation between motor movements and the PPC may reflect progressive coordination of activity in PPC with sensorimotor inputs from cortex and thalamus, regulating varying aspects of associational connectivity at different developmental stages before the start of memory directed spatial navigation.

Our fMOI recordings permitted a global network analysis of activity at unprecedented temporal and spatial scale across the cortex. Since the current work focused on recordings of ongoing cortical activity in the absence of directed sensory stimulation or task engagement, our results are comparable to studies of functional connectivity using resting state fMRI^44^ in humans or animal models. Cortical activity exhibited spatial and temporal features that were consistent between animals and across ages. Most strikingly, much of the activity was symmetrical across the two hemispheres, similar to the interhemispheric resting state connectivity between homotopic cortical regions observed in human fetal^45^ or infant^46^ resting state fMRI. Large scale brain networks share features of ‘small-world’ network organization^47^, such as short average path lengths (efficient global information transfer like a random network) and high clustering among neighboring nodes (robustness to random error like a lattice network)^37^, along with the presence of high network modularity (presence of community subnetworks) and hubs (nodes having high degree of connections and centrality within or between communities)^48,49^. Our network analysis showed that the developing functional network in rodent cortex has a short average path length with high clustering and a significant small-world index^50^ (**Supplementary Fig. 4**), consistent with reports of small-world properties in human resting state networks at birth^50,51^. We found that the developing mouse brain network is comprised of 3–5 distinct functional modules among cortical areas, reminiscent of the 4 functional networks in neocortex observed with resting state fMRI in the fetal, infant, and adult human brain^50,52,53^. Furthermore, our data revealed several hub areas, including secondary motor cortex (M2, cingulate, mPFC), posterior parietal cortex (PPC), and retrosplenial cortex (RSA), that are remarkably consistent with homolgous cortical hub regions identified in resting state fMRI networks in human^49,54^ (mPFC, anterior cingulate, lateral parietal cortex, posterior cingulate cortex) and adult anesthetised rat^55^ (mPFC, cingulate, PPC, RSA). Some hub regions, such as M2 and PPC, shifted module memberships during development, perhaps signaling their importance as associative areas and connector hubs between communities, which is comparable to developmental changes in module membership in human resting state fMRI cortical networks^48,52^. The increasing strength of intra- and inter-hemispheric connections with age is consistent with greater long-range connection strength between disparate brain regions during brain development^48,49^. This work shows that properties of functional network architecture emerge and may be shaped very early in cortical development. Given that pathophysiological changes in functional connectivity of large scale brain networks is thought to be a key feature of a number of neurological disorders, including autism and schizophrenia^56,57^, extension of these fMOI methods to genetic and environmental disease models could help in understanding altered network dynamics that underlie neurodevelopmental disorders.

There are several advantages of our fMOI approach for assessing cortical activity dynamics in animal models compared with other techniques for recording population activity in the brain. One is the combination of high temporal and spatial resolution relative to techniques like functional magnetic resonance imaging (fMRI), intrinsic signal imaging or traditional EEG and electrode recordings. A second advantage is that fMOI provides a relatively non-invasive method of recording ongoing neuronal population activity in mice transcranially, without confounds associated with anesthesia, compared with traditional electrode recordings, voltage-sensitive dye-, intrinsic signal-, or two-photon imaging based recordings. Another benefit is that the calcium dynamics imaged with fMOI provide a direct readout of ongoing neuronal activity, and signals can be localized by select expression of the genetic calcium indicator in distinct cell populations. Furthermore, the isolation of functional response from movement artifacts is relatively simple with fMOI in comparison to many population recording methods. Finally, the spatial scale of the fMOI approach surpasses that of traditional techniques, with nearly complete coverage of the neocortex, enabling simultaneous recording of numerous sensory-motor areas throughout the hemispheres. Given the notable parallels we’ve demonstrated here between the functional development of mouse and human cortical network architectures together with the broad scope and high spatiotemporal resolution of fMOI for direct recordings of neuronal activity, extension of this work may prove useful in understanding the nature of fMRI connectivity and its link to ongoing brain activity patterns.

This study demonstrates that ongoing activity in developing cortex is not random – it is coordinated in space and time across the entire neocortex. These structured whole brain activity patterns may play key roles in the activity-dependent development of local and global circuit connectivity throughout the nervous system. Moreover, fMOI provides a new, non-invasive method for studying functional connectivity in animal models, and our results demonstrate how the functional cortical network architecture in developing mouse shares common fundamental features with human resting state networks. The simplicity and robustness of functional mesoscale optical imaging will likely be key to integrative assessments of altered brain activity dynamics in animal models for neurological disorders.

## Methods Summary

Rx-Cre:GCaMP3 (Ai38) or SNAP25-GCaMP6 (Ai103) mice aged between postnatal day 2 to 13 (P2–P13) were prepared for transcranial optical imaging as described previously^17^. Calcium imaging was performed in vivo using wide-field epifluoresence microscopy with a 1x macro objective and a pco.edge sCMOS camera after a 1-hour recovery period from general anesthesia. Automated image segmentation and calcium event detection was performed using custom, freely available MATLAB routines.

### Acknowledgements

We thank Y. Zhang for technical support. We thank C. Messinger for help with initial SERT-Cre:tdTomato^f/f^ histology. We would like to thank Emily Finn, Todd Constable, and Daeyeol Lee for valuable comments on the manuscript. This work was supported by NIH Grants RR19895, RR029676–01 for the Yale University Biomedical High Performance Computing Center and NIH grants P30 EY000785, R01 EY015788 to M.C.C. M.C.C. also thanks the family of William Ziegler III for their support.

### Author Contributions

J.B.A. and M.C.C. designed the experiments. J.B.A. performed in vivo imaging experiments, wrote the image processing and data analysis routines, and analyzed the recordings. H.Z. created the GCaMP3 and GCaMP6 mouse lines. J.B.A. and M.C.C. wrote the manuscript.

## Methods

### Animals

Animal care and use was performed in compliance with the Yale IACUC, U. S. Department of Health and Human Services and Institution guidelines. Neonatal Ai38 floxed GCaMP3 reporter mice (JAX no. 014538)^58^ crossed with *Rx-Cre*^59^ or SNAP25-GCaMP6 (Ai103) transgenic mice aged 2–13 days (P2–P13) after birth (P0) were used.

### Transgenic mice generation

The *Snap25* locus was targeted with homologous recombination to insert LSL-F2A-GFP at the endogenous stop codon, using a targeting vector containing the following components: 5′ arm – F3 – last ∼300bp of intron 7 – exon 8 up to the endogenous stop codon – loxP – stop codons – PGK polyA – loxP – F2A-EGFP – WPRE – bGH polyA – AttB – pPGK – neomycin-resistant gene – PGKpA – F5 – mRNA splice acceptor – domain 2 from the hygromycin-resistant gene – SV40 polyA – AttP – 3’ arm. Targeting constructs were generated using a combination of molecular cloning, gene synthesis (GenScript, Piscataway, US) and Red/ET recombineering (Gene Bridges, Heidelberg, DE). The 129S6B6F1 ES cell line, G4, was used for the gene targeting. Correctly targeted neomycin resistant clones were identified by PCR, and then confirmed by Southern blot. One ES clone that gave high percentage chimeras following blastocyst injection was used in a Flp recombinase-mediated cassette exchange (RMCE) transfection to switch the LSL-F2A-GFP expression unit for a T2A-GCaMP6s expression unit. The replacement vector included: F3 – last ∼250bp of intron 7 – exon 8 up to the endogenous stop codon – T2A-GCaMP6s – WPRE – bGH polyA – AttB – pPGK – domain 1 from the hygromycin-resistant gene – mRNA splice donor – F5. Following co-transfection with a pCAG-Flpe plasmid, hygromycin resistant colonies were screened for correct 5′ and 3′ junctions and for lack of the original vector sequence. Correct clones were used in blastocyst injections to obtain germline transmission. Resulting mice were crossed to the Rosa26-PhiC31 mice (JAX Stock # 007743) to delete the pPGK-hygro selection marker cassette (in between the AttB and AttP sites), and then backcrossed to C57BL/6J mice and maintained in C57BL/6J congenic background.

### Surgical procedure for in vivo imaging

Mice aged P2–P13 were deeply anesthetized with isoflurane (2.5%) in oxygen and then placed on a heating pad set to 35°C via a isothermic temperature monitor (NPI TC–20, ALA Scientific). Local anesthesia was produced by subcutaneous injection (0.05 ml) of 1% Xylocaine (10 mg/ml lidocaine/0.01 mg/ml epinephrine, AstraZeneca) under the scalp. After removal of the scalp, steel head posts were fixed to the exposed skull using cyanoacrylate glue. A 1 hr recovery period in the dark under continuously delivered medical oxygen with isoflurane at 0% was allowed after surgical preparation and the mouse was surrounded by a cotton ball nest. This recovery period was the typical minimum time required for spontaneous waves of activity to develop in the visual system after the cessation of deep anesthesia^17^. A red LED (Radioshack) and photodiode sampled at 25 kHz with a Power1401 (Cambridge Electronic Design) were positioned to monitor respiratory rate and limb/body movements.

### Wide field calcium imaging

A 16 bit CMOS camera (pco.edge, PCO) coupled to a Zeiss AxioZoom V16 microscope with 1X macro objective was used to image transcranial calcium dynamics. Epifluorescent illumination was provided by a DC stabilized Hg^2+^ light source (X-Cite, EXFO) through a EGFP filter cube set (Zeiss) with the minimum illumination intensity that gave detectable calcium signals using a exposure of 200 msec. Image frames corresponding to a field of view of 6 × 8 mm, 10 × 12 mm, or 20 × 24 mm were acquired at a rate of 5 or 10 frames per second (200 ms or 100 ms frame period). Each recording consisted of a single, continuously acquired movie for 10 min.

### Calcium signal detection

Image processing and calcium signal detection was performed using custom software (available at http://github.com/ackman678/wholeBrainDX) written in MATLAB (The Mathworks, Natick, MA). The mean pixel intensity at each pixel location, *F0* was subtracted and normalized to each frame, *Ft* of the movie to form a dF/F array: *A* = (*Ft* − *F0*)/*F0*. A background estimate was calculated and subtracted from every frame of *A* with a top hat filter using a disk shaped structuring element with radius of 620 μm. Each frame was smoothed with a Gaussian having a standard deviation of 56 μm and a signal intensity threshold, *T* was computed using Otsu’s method on the set of histogram of pixel intensities in *A* that corresponded to the set of pixels at the 99th percentile from the Sobel gradient transformation of *A*. Calcium domain signals were automatically segmented as contiguously connected components in space and time from the binary mask array, *A* > *T*. Components located outside the cortical hemisphere boundaries or having an area < 50 pixels or a duration of 1 frame were ignored.

### Statistical analysis

Data sets were analyzed and plotted using custom routines implemented in MATLAB (The Mathworks, Natick, MA) and in R (The R Project for Statistical Computing, http://www.r-project.org) with the ggplot2 plotting library (http://ggplot2.org). Distribution means were compared using two-sample Student’s t-tests unless otherwise noted (Wilcoxon Rank Sum test was used for small, non-normally distributed data sets) or using ANOVA followed by Tukey’s HSD post-hoc test when analyzing the effects of multiple grouping factors (p < 0.05 set as significance). Values are reported as means with the standard error of the mean or medians with the median absolute deviation unless otherwise noted. All boxplots report the median, the 25^th^ and 75^th^ percentiles as lower and upper box hinges (1^st^ and 3^rd^ quartiles), the data range as lower and upper whiskers (lowest and highest data values within 1.5 ∗ IQR of the lower and upper hinge, where IQR is the difference between the 3^rd^ and 1^st^ quartiles), and outliers as individual gray points (data values beyond the whisker range).

### Calcium domain analysis

The mean width in the medial-lateral and height in the rostral-caudal dimensions of the bounding box fitted to each segmented calcium domain signal was taken to be the domain diameter. The number of contiguous frames (bounding box depth) for each segmented calcium domain was taken to be the domain duration. The mean and maximum pixel intensities from *A* within each domain were taken as the mean and maximum domain amplitudes. After functional parcellation of the cortical hemipsheres (Fig. 1; Supplementary Fig. 1), domains were assigned areal membership by intersection of the domain centroid with a cortical area’s pixel mask. The number of individual domains per recording within a hemisphere or cortical area was taken to be domain frequency.

### Wave motion analysis

Optical flow was computed using the Lucas-Kanade method on the binary movie array from all the segmented calcium domain masks for a recording. The motion magnitude, *R* was the velocity vector sum for each domain. The wave motion index for each domain was calculated as *R*^2^/*D*^2^, where *D* was the domain diameter.

### Motor movement analysis

A binary movie array from all the segmented calcium domain masks for a recording was intersected with masks representing different cortical area parcellations. The total number of active pixels per frame expressed as a fraction of the total number of possible pixels per frame for each cortical area gave active pixel fraction time courses for each cortical area in each recording. Movement signals acquired with the photodiode were bandpass filtered using an 8-order, 1-20 Hz pass band elliptic filter, and then rectified and downsampled to the movie frame rate to give a movement time course signal that corresponded to displacements of the limbs and body excluding those from respiration. Cross-correlations between combinations of cortical active fraction time courses and the movement signal time course were computed, and the Pearson’s correlation coefficient at the zeroth time lag was obtained.

### Hemisphere correlation analysis

A binary movie array from all the segmented calcium domain masks for a recording was intersected with masks representing the cortical hemispheres. The total number of active pixels per frame expressed as a fraction of the total number of possible pixels per frame for each cortical area gave active pixel fraction time courses for each hemisphere in each recording. The mean, normalized medial-lateral and anterior-posterior positions for all active pixels within each hemisphere during coactive frames gave spatial center of mass timecourses for each recording. Pearson’s correlation coefficient was calculated between the active pixel fraction time courses and the activity center of mass time courses to give the temporal and spatial correlation for each movie.

### Functional connectivity analysis

A binary movie array from all the segmented calcium domain masks for a recording was intersected with masks representing different cortical area parcellations. The total number of active pixels per frame expressed as a fraction of the total number of possible pixels per frame for each cortical area gave active pixel fraction time courses for each cortical area in each recording. Correlation matrices were calculated for each recording by computing pairwise Pearson’s correlation coefficients, *r*, from the matrix containing the cortical active pixel fraction time courses. The mean correlation matrix for each age group was computed and then the binarized correlation matrix at *r* > 0.15 (the maximum *r* value where the 3 largest communities were connected in the graphs at all age groups) was used to form an adjacency matrix with each node representing a cortical area and each edge representing an association between a pair of nodes at weight, *r.*

### Network analysis

Graph theoretical analyses were performed using the igraph network analysis software library (http://igraph.org). Community structure was detected within each functional association matrix using a greedy optimization algorithm that maximizes the graph modularity score to perform hierarchical clustering^36,60^, where the modularity score measures the fraction of edges within modules for a graph partition compared with that of a randomized equivalent network. Network graphs were plotted using an anatomical layout or using a force-directed graph layout^61^ with nodes colored by module membership and edges connecting nodes reflecting the edge weight, *r*. Node degree was the number of connections that link a vertex to the rest of the network. The average path length, *L* of a graph was the mean of the shortest paths (fewest number of edges) between all pairs of nodes. The random average path length, *L*_*r*_ was the mean of the shortest paths in a set of 1000 equivalent random networks that had the same degree sequence as the original graph. The local clustering coefficient was the ratio of the triangles connected to the node and the triples centered on the node, measuring the probability that two neighbors of a node are also connected. The global clustering coefficient, *C* was the ratio of the triangles and connected triples in the graph. The random global clustering coefficient, *C*_*r*_ was the mean of the clustering coefficients in a set of 1000 equivalent random networks that had the same degree sequence as the original graph. The small-world index was calculated as the ratio of the normalized clustering coefficient (*C*/*C*_*r*_) and the normalized path length (*L*/*L*_*r*_), where a small-world index > 1 indicates a small-world network organization^50,62^. Node strength was the column sums in the weighted adjacency matrix. Betweenness centrality scores corresponded to the fraction of all shortest paths that pass through a node^63^. Eigenvector centrality scores were the values of the first eigenvector of the association matrix^64,65^, reflecting for each node the sum of direct and indirect connections of every length in a network.

